# Towards automation of germline variant curation in clinical cancer genetics

**DOI:** 10.1101/295865

**Authors:** Vignesh Ravichandran, Zarina Shameer, Yelena Kernel, Michael Walsh, Karen Cadoo, Steven Lipkin, Diana Mandelker, Liying Zhang, Zsofia Stadler, Mark Robson, Kenneth Offit, Joseph Vijai

## Abstract

Cancer care professionals are confronted with interpreting results from multiplexed gene sequencing of patients at hereditary risk for cancer. Assessments for variant classification now require orthogonal data searches, requiring aggregation of multiple lines of evidence from diverse resources. The burden of evidence for each variant to meet thresholds for pathogenicity or actionability now poses a growing challenge for those seeking to counsel patients and families following germline genetic testing. A computational algorithm that automates, provides uniformity and significantly accelerates this interpretive process is needed. The tool described here, Pathogenicity of Mutation Analyzer (PathoMAN) automates germline genomic variant curation from clinical sequencing based on ACMG guidelines. PathoMAN aggregates multiple tracks of genomic, protein and disease specific information from public sources. We compared expert manually curated variant data from studies on (i) prostate cancer (ii) breast cancer and (iii) ClinVar to assess performance. PathoMAN achieves high concordance (83.1% pathogenic, 75.5% benign) and negligible discordance (0.04% pathogenic, 0.9% benign) when contrasted against expert curation. Some loss of resolution (8.6% pathogenic, 23.64% benign) and gain of resolution (6.6% pathogenic, 1.6% benign) was also observed. We highlight the advantages and weaknesses related to the programmable automation of variant classification. We also propose a new nosology for the five ACMG classes to facilitate more accurate reporting to ClinVar. The proposed refinements will enhance utility of ClinVar to allow further automation in cancer genetics. PathoMAN will reduce the manual workload of domain level experts. It provides a substantial advance in rapid classification of genetic variants by generating robust models using a knowledge-base of diverse genetic data https://pathoman.mskcc.org.

## Introduction

The uptake of genetic testing and targeted resequencing of cancer susceptibility genes to facilitate precision cancer prevention and early diagnosis has grown exponentially in concert with the decreasing costs of next generation sequencing (NGS)^1;2^. A major challenge is the interpretation of sequence variants which result from clinical sequencing. American College of Medical Genetics and Genomics (ACMG) and the Association of Molecular Pathology (AMP) have published guidelines on interpretation of germline variants taking into account not only their pathogenicity but also their clinical actionability^3^. Nonetheless, germline variant classification continues to pose an immense burden on the time and resources of diagnostic molecular labs and cancer care professionals. The ACMG classification schema requires manually exploring multiple lines of public data, other orthogonal data sources and from the literature; then aggregation and scoring to provide evidence for classifying variants^4^. There is currently no automated computational framework for classifying genetic variants that is widely available. We developed PathoMAN, a computational resource that automates this classification with uniformity, transparency and speed, in order to facilitate variant curation for the cancer genetics community.

PathoMAN is a Python based variant curation algorithm that classifies germline genomic variants that are detected in the context of clinical cancer genetics sequencing. The schema is inspired by ACMG/AMP classification^3^. It aggregates multiple tracks of genetic and molecular evidences using variant annotators and from public repositories containing evidence for pathogenicity assertion. The compiled data is then used in 28 distinct categories addressing the type of the mutation, its biological impact, in silico predictions, presence in the control cohort as well as the familial information and inheritance pattern of the mutation. The aggregate score resulting from these categories is used in classifying the variant as Pathogenic (P), Likely-Pathogenic (LP), Benign (B), Likely-Benign (LB) or Variant of Uncertain Significance (VUS).

The performance of PathoMAN was measured by re-evaluating expertly curated germline cancer variants in cancer genes compiled from two large published studies on breast and prostate cancer. In this study, we also assess the frequency of clinically actionable mutations present in general population using ExAC data in cancer susceptible and cancer predisposing genes as well as in ClinVar. We test the application of ACMG criteria for germline cancer variants and address the bottleneck of variant curation in using automated algorithms in variant classification.

## Materials and Methods

### Test datasets

We tested 11,196 germline variants in 76 genes of highest cancer relevance to describe and compare performance of the algorithm in cancer related genomic variant data. Clinically actionable variants (P/LP) in these 76 genes are reported back to patients at MSKCC as part of the IMPACT^®^ assay using the appropriate IRB approved protocol^5;6^. Several of the genes in this set are strong cancer predisposition genes, while others are putative candidate genes across one or more syndromic cancers. We have selected this gene list, referred as IMPACT-76^7^,(**Suppl Table 1**) uniformly throughout this manuscript. We chose exonic and essential splice site (+/− 1,2) variants in the IMPACT-76 genes from three manually curated datasets - (i) prostate cancer study^8^; (ii) breast cancer study^9^ and (iii) ClinVar^10;11^. The germline variants in the cancer datasets were curated by experts using the ACMG classification guidelines. Some adjustments or manual overrides were performed based on the interpretation of the variant in a disease and literature evidences^9^. It should be also be noted, that in the breast dataset analysis, the exome sequencing project (EVS6500) and 1000 genomes for population allele frequencies and the ClinVar version available in early 2016 were used.

The Exome Aggregation Consortium^12^(ExAC) is a joint effort to aggregate exome sequencing data from fourteen large sequencing projects to provide summary data such as ethnicity specific allele frequency for a wider scientific community. The ExAC-noTCGA data is a subset of 53,105 samples and it doesn’t include 7601 The Cancer Genome Atlas (TCGA) cancer germline samples. The variants in test data were compared against ExAC-noTCGA Non-Finnish European population and were considered not to have *de novo* evidence and co-segregation, as this information was unavailable for these datasets. However, PathoMAN can use such information, if available to assign ACMG classes. The results were compared against the manual curation. They are reported here in four categories: concordance, discordance, loss of resolution and gain of resolution.

When the reported P/LP and B/LB variants are re-classified as P/LP and B/LB respectively by PathoMAN, then the results are considered concordant. Similarly when reported P/LP and B/LB variants are re-classified as B/LB and P/LP by PathoMAN respectively, then the variants are considered discordant. When reported P/LP or B/LB variants are re-classified as VUS by PathoMAN, then they are placed in the loss of resolution (LOR) category and when the reported VUS are re-classified as P/LP or B/LB, then are considered as gain of resolution (GOR) category (**Table 1**). As we describe below, these evaluations aid in understanding the real-world usage of the eight ACMG categories of evidence in cancer genomics, and in their ability to discriminate between P/LP, B/LB and VUS.

**Table 1:** Rating the comparison of results between PathoMAN variant curation and domain expert-curation of germline variants.

| Rating | Reported | PathoMAN |
| --- | --- | --- |
| Concordance | P/LP; B/LB | P/LP; B/LB |
| Discordance | P/LP; B/LB | B/LB; P/LP |
| Loss of Resolution (LOR) | P/LP; B/LB | VUS |
| Gain of Resolution (GOR) | VUS | P/LP; B/LB |

P-pathogenic, LP-likely pathogenic; B-benign; LB-likely benign; VUS-Variants of Uncertain Significance.

### ExAC subset of IMPACT-76

The ExAC^12^ is a public resource often utilized as convenience controls for several human traits in case-control studies. We wanted to estimate the burden of variants in ExAC-noTCGA as classified by PathoMAN and contrast against known information in ClinVar. We selected 55,566 variants from IMPACT-76 genes which were in exonic or essential splice site regions. This is not considered part of the test datasets descibed earlier, as ExAC data is used as part of the ACMG criteria PS4, PM2, BA1, BS1 and BS2.

### ACMG/AMP guidelines

The 2015 ACMG/AMP guidelines consist of 16 criteria that aid in classifiying pathogenicity and 12 criteria that aid for benignity. Pathogenicity criteria were broadly grouped as: very strong evidence (PVS1), strong evidence (PS1-PS4), moderate evidence (PM1 – PM5) and supporting evidence (PP1-PP5). Benign criteria are broadly grouped as: standalone evidence (BA1), strong evidence (BS1-BS4) and supporting evidence (BP1-BP7). This classification system resolved a variant as pathogenic or benign based on eight components – population frequency data, genomic annotation and computational predictive data, functional data, segregation data, *de novo* data, allelic/genotypic data, public databases and literature and other data (Table 2). The variant classification criteria used by PathoMAN were inspired by these ACMG/AMP guidelines, although they do not precisely adhere to these published norms. We describe below the criteria and their modifications for variant classification by PathoMAN for cancer genetics.

### Variant Annotation

PathoMAN makes use of CAVA^13^ for genomic annotation, dbNSFP^14–16^ using Annovar^17^ for in silico predictions, ExAC-noTCGA and gnomAD^12^ for public control frequencies, ClinVar^10,11^ for public evidences, and curated list of variants from the literature for functional evidences.

### Determination of PVS1: null variants

Curated lists of cancer-causing genes from the literature, various genetic testing panels,^18–22^ and OMIM genes that causes autosomal dominant disease^23^ were aggregated. If the variant in a gene from this list was a Tier 1 mutation (frameshift, truncating, essential splice variant and initiation codon), and not present in thelast exon, then PVS1 was scored 1. A gene with a functional domain encoded by the last exon, such as *ATM*, was an exception to the last exon criteria^24^. PVS1 was not scored for *BRCA2* mutations observed after the polymorphic stop rs11571833 (K3326X).

**Table 2:**
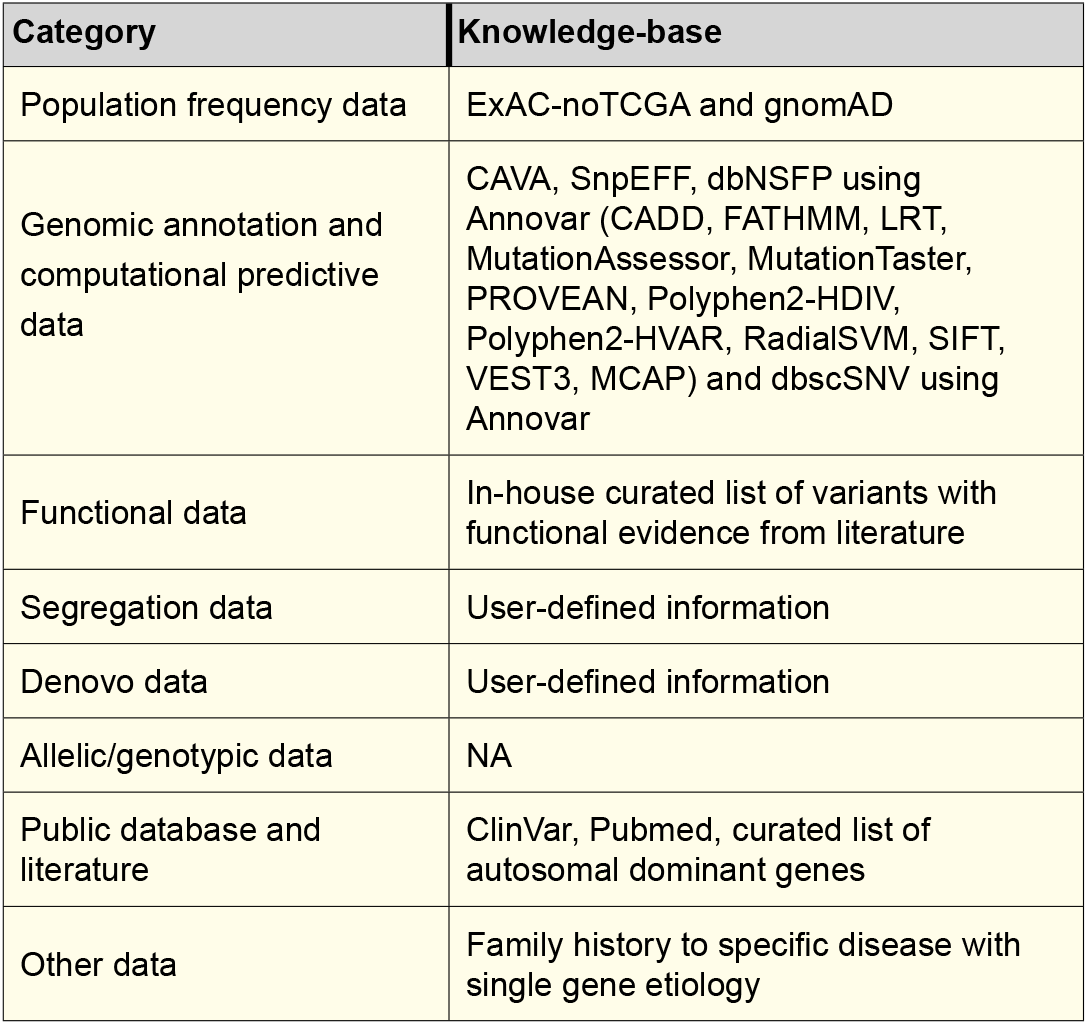
List of data sources and annotators used in building PathoMAN’s knowledge-base

### Determination of PS1 and PM5: known pathogenic missense

If a missense variant was reported as pathogenic by multiple submitters with no conflicts and had a gold star of 2 or more in ClinVar^25,10,11^, irrespective of the alternative allele but leading to the same amino acid change, then PS1 was scored 1. If a missense variant was not seen in ClinVar but had another pathogenic missense variant at the same amino acid with a different amino acid change, then PM5 was coded 1.

### Determination of PS3 or BS3: strong prior evidence of pathogenic or benign

An aggregated select list of reported pathogenic and benign variants from the literature^2;26-29^ were used as a knowledge-base for PS3/BS3. If the variants were in the curated list (missense variants in BRCA1/2 reported by ENIGMA), or if it was a truncating variant and ClinVar had reported it as pathogenic or benign with a gold star 2 or more, then PS3 or BS3 was coded 1. Our selective use of ClinVar assigns higher confidence for the truncating variants and select missense variants reported by domain experts that are either pathogenic or benign. We also include published saturation editing experimental evidence for *BRCA1*.

### Determination of PS4 PM2 BA1 BS1 and BS2: rarity and enrichment of variant in cases

If a variant was present in aggregated public controls such as the ExAC-noTCGA dataset and gnomAD^12^ with an allele frequency greater than 5%, then the variant was coded 1 for BA1. If the variant had an allele frequency in public controls between 1% and 5% then BS1 was coded 1. If the variant was also present in a homozygous form in the public controls, then BS2 was coded 1. If the variant was absent from ExAC-noTCGA data or gnomAD general population data, and then PM2 was coded 1. For variants, not scored as BA1, BS1, BS2 or PM2, Fishers Exact test was performed against the user defined population (ExAC-noTCGA or gnomAD). The population included all the major groups (NFE, FIN, SAS, AMR, AFR, EAS) in ExAC and ASJ population in the gnomAD database. If the odds ratio was greater than 3 and p-value less than 0.05, then the variant was given a score of 1 for PS4. This was a robust measure for weighting pathogenicity in uncommon variants and non-singletons.

### Determination of PM1: membership in a protein domain of functional significance

If the amino acid that was being altered by the mutation was present in a protein domain, or a residue involved in signalling, binding with other proteins, or in an active site, then PM1 was coded 1. Currently, we use Uniprot for annotation of protein features^30^. We acknowledge the incremental value of a curated somatic hotspot list (http://cancerhotspots.org) to aid in this classification.

### Determination of PM4 BP3: genomic complexity and context of the variant

If the mutation was an in-frame insertion/deletion or a stop loss in a non-repetitive region, then the variant was coded 1 for PM4. Instead, if it was an in-frame insertion/deletion in a repetitive region, then BP3 was coded 1. The repeat masker track from UCSC genome browser was used for this^31–34^ criteria.

### Determination of PP3 BP4: In silico prediction of deleteriousness

We used Annovar17to annotate the variants with dbNSFP^14-16,35^ track to get results of deleteriousness predictions from 12 in silico algorithms – CADD^36^, FATHMM^37^, LRT^38^, MutationAssessor^39^, MutationTaster^40^, PROVEAN^41^, Polyphen2-HDIV^42^, Polyphen2-HVAR^42^, RadialSVM^35^, SIFT^43^, VEST3^44^and M-CAP^45^. Use of an ensemble of in silico prediction algorithms improves prediction across a wide range of genes and cancer types^46^. Hence if more than 7 (>50%) algorithms call a variant deleterious, then the variant was coded 1 for PP3. Otherwise, BP4 was coded 1. In contrast, many of the old ClinVar records relied on only SIFT or Polyphen.

### Determination of PP5 BP6: Known variant with insufficient details

If the variant was in ClinVar^10,11^ with gold star less than 2 and was pathogenic, then PP5 was scored 1. If it’s benign, BP6 was scored 1. Thus, a variant reported once in ClinVar does not command high value in the pathogenicity determination, but can be upgraded depending on other ancillary information tagged to it.

### Determination of BP7: synonymous variants

For synonymous silent mutations, we used adaptive boosting and random forest scores from dbscSNV^47^, which if it was less than 0.6, then BP7 was scored 1. dbscSNV is a database of precomputed prediction scores for SNVs, that may occur in splice consensus regions. Higher scores reflect the variants effect in splicing^47^.

### Determination of PP2 and BP1: missense driven disease genes

We used ClinVar^10; 11^ to collect all reported missense variants per gene. We then selected the confident (gold star 2 or more) pathogenic and benign calls. The list of genes with higher ratio of pathogenic to benign variants called missense-driven pathogenic genes and the list of genes with lower ratio of pathogenic to benign variants were called missense-driven benign genes. Any missense variant in the missense-driven pathogenic gene list was scored PP2 and any missense variant in the missense -driven benign gene list was scored BP1. Classic example of such genes are PTEN^49^ and *TP53*^50^. Almost all *PTEN* missense mutations were pathogenic (**Supp Figure 1**).

### Determination of PS2, PM6, PP1 and BS4: denovo and cosegregation

ACMG criteria require both paternity and maternity confirmed for *de novo* variants. PathoMAN requires user input for *de novo* status and segregation information for classification. Three options for *de novo* evidence includes, *de novo* with both paternity and maternity confirmed, *de novo* without paternity or maternity confirmed and no *de novo* evidence at al. Similarly, for cosegregation evidence, the options provided were co-segregation with the disease, lack of co-segregation with the disease and no co-segregation. For larger trio studies, in the future, we expect to include a module that looks for *de novo* variants computationally from a mutisample VCF and pedigree file information.

### Determination of PM3, BP2: recessive inheritance

These two categories apply to variants with a recessive disease. Germline cancer variants are generally associated with cancer predisposition syndromes in an autosomal dominant inheritance pattern. Hence, PM3 and BP2 doesn’t apply for current PathoMAN variant classification. They were scored 0. We expect to add compound heterozygosity to the next version of the algorithm. We also will incorporate select gene variants in mismatch repair genes that can be classified by this category into the knowledge-base.

### Determination of PP4, BP5: disease specific conditions

PP4 and BP5, per ACMG were two criteria to consider in a patient’s predisposition to a specific disease with single gene aetiology. Cancer is a disease with multi-gene aetiology although certain genes such as *RB1* may be strong candidates for PP4. Once we incorporate a pedigree file, and cancer phenotype variables, we should be able to apply these criteria. In the current iteration, these were scored 0.

The final classification schema was based on the original ACMG scoring and pathogenicity was predicted for the datasets described.

### Usage of ClinVar (Release 12/28/2017) in PathoMAN

ClinVar is the most popular database that processes submissions of human DNA variants and assertions made regarding their clinical significance. One of its stated goals is to support computational re-evaluation of genotypes and assertions. ClinVar currently does not provide detailed information on pathogenicity assertions, unlike another initiative ClinGen^51; 52^, which aims to build a more discriminatory and accurate assessment of variant pathogenicity. However, access to the ClinGen interface is at the moment restricted to a smaller niche group. PathoMAN uses ClinVar as one of its knowledge-base. We use the ClinVar data parser tool^53^ to extract important fields that will aid in variant classification. PathoMAN takes advantage of the gold star system awarded to variants with highest evidences supporting the assertion of clinical significance. PathoMAN up-weights variants with gold star 2 or more in the knowledge-base. We selected variants with gold star < 2 and with the term “cancer” in traits field and used them as another test dataset. This test data set was filtered for only variants in the exonic and essential splice regions in IMPACT-76^54^ genes (n=9164) variants (Supp Table 1).

## Results

### PathoMAN versus manual curation for test datasets

PathoMAN is a variant classification algorithm, provided as a web based service, that either allows user to query single variants or upload a file with a batch of variants. The web-tool is created using Flask web framework. Single variant query works on chromosome, position, reference allele, alternative allele, allele count, allele number, *de novo* status, co-segregation status and preferred control population sub-group. Batch upload requires six columns that follow VCF 4.2 format [*chr, pos, ref, alt, ac, an*]. User can select *de novo* status, co-segregation status and preferred control population sub group. Our pipeline is currently equipped to annotate a minimal VCF and prepare it for PathoMAN variant classification. The result of a single variant query is displayed back on to the web-page while the batch upload results will be mailed back to the registered user. For an annotated VCF file containing 1000 variants, PathoMAN takes 6 minutes, which is 3.6 seconds per variant. This provides a massive advantage in terms of speed, uniformity, efficiency and service assurance than manual curation.

Overall the test dataset contained 11,196 variants in 76 genes with reported expert curation from three groups–(i) prostate cancer study^8^; (ii) breast cancer study^9^and (iii) ClinVar^10,11^. Missense mutations account for 71% of the variants, Frameshift variants 14%, Stop gain variants 5% and the rest distributed among splice variants, in-frame insertion/deletion and stop loss in the test dataset (**Figure 1A, Table 3**). We annotated the variants with CAVA, Annovar, ExAC noTCGA, gnomAD and ClinVar and prepared the VCFs for PathoMAN.

**Figure 1A:**
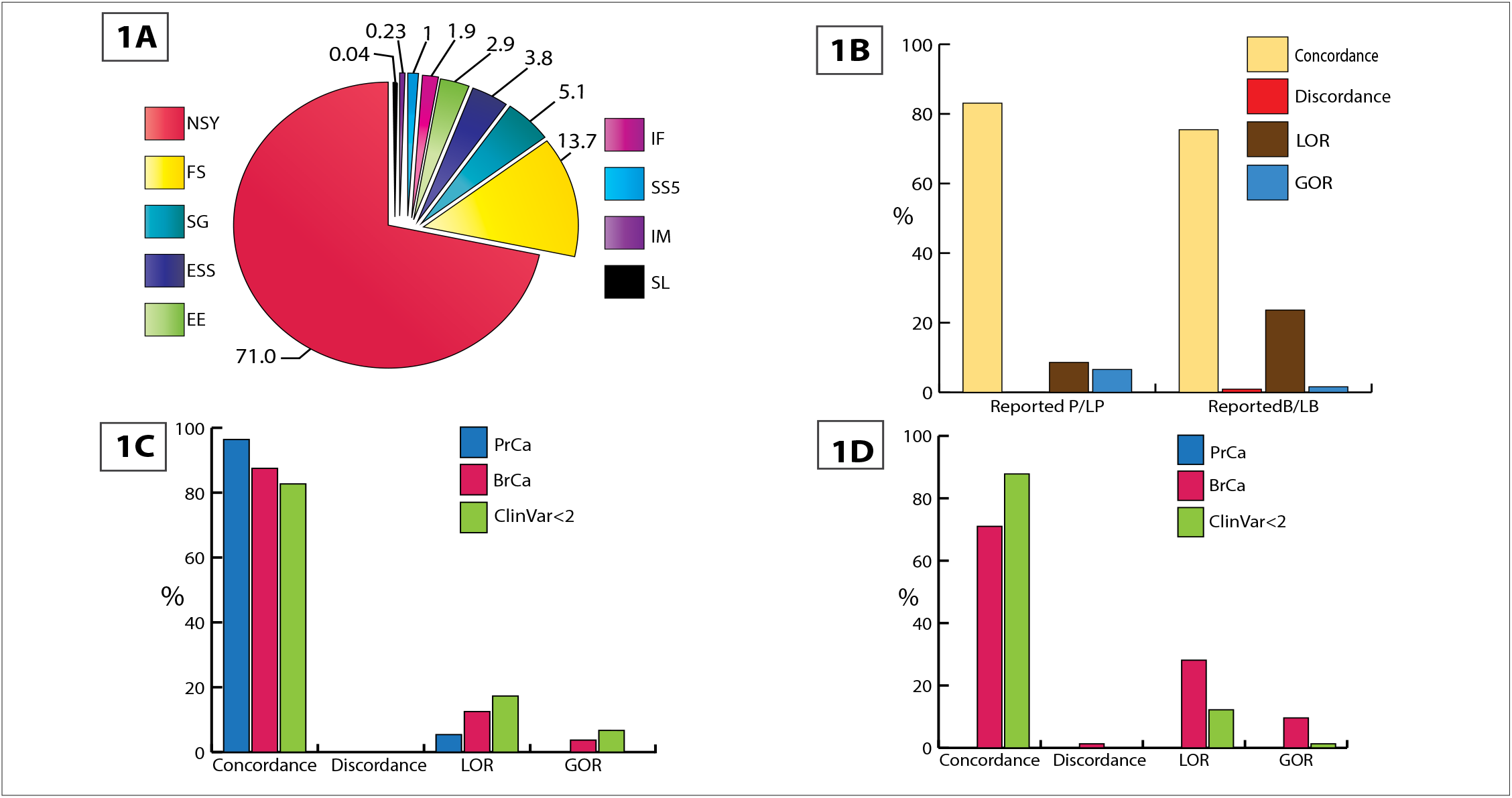
Distribution of mutations in test data by variant class. (Stop loss - SL; Start codon alteration - IM; Splice variant that alters +5 splice site - SS5; Inframe ins/del - IF; Alters first 3 bases of codon - EE; Essential splice site +/− 1, 2 - ESS; Stop gain - SG; Frameshift - FS; Non-synonymous - NSY). **Figure 1B: Overall comparison results of PathoMAN variant curation and expert curation for germline variants from test dataset. Figure 1C: PathoMAN results for reported pathogenic and likely pathogenic germline variants in test dataset. Figure 1D: PathoMAN results for reported benign and likely benign germline variants in test dataset**.

PathoMAN achieves an overall concordance of 83.1% for P/LP variants and 75.5% for B/LB variants. A minimal discordance of 0.04% (P/LP) and 0.9% (B/LB) and a LOR of 8.6% (P/LP) and 23.6% (B/LB) were observed. PathoMAN achieves GOR by resolving 6.6% of the VUS as P/LP and 1.59% as B/LB (**Figure 1B-D**). We estimated reliability of recall using the Cohen’s kappa coefficient (*K*) between the manual curation and PathoMAN classification for the test dataset. The number of observed agreements was 89.2% (9988 unique variants) (*K* = 0.74, CI 95% 0.73-0.75).

Out of the 2,535 P/LP variants reported by manual curation, PathoMAN showed very high concordance for frameshift, essential splice sites and truncating variants. PathoMAN failed to resolve reported P/LP for 233 missense variants (**Table 4A**). All of these were low confidence variants in the ClinVar dataset with no assertion criteria provided (gold star=0) or variants with a single submitter (gold star=1). Similarly for reported benign variants (n=440), we observe high concordance with the exception of missense variant class. LOR is observed for 90 missense variants with conflicting reports in ClinVar (**Table 4B**). Many rare variants that are reported from clinical sequencing/studies have no public records documenting their pathogenicity classification and there may be insufficient evidence to call them either pathogenic or benign. These variants have been called VUS by the manual curators (n=8221 variants) at the time. PathoMAN is able to resolve 6.7% of the VUS as LP and 1.6% as LB (**Table 4C**).

### PathoMAN results for ExAC (no TCGA) dataset

We asked if we could predict the different classes of ACMG mutations in this public resource using PathoMAN. We selected 55566 exonic and essential splice variants from IMPACT-76 genes and classified them. Overall, PathoMAN calls <1% of the heterozygous genotypes in ExAC noTCGA dataset, from 53,105 samples, as P/LP. Further we tabulated pathogenic variant burden by genes and compared them against ClinVar (**Table 5**). The results show, PathoMAN calls a similar number of variants reported in ClinVar for the bona_fide cancer genes like *BRCA1* and *BRCA2*. PathoMAN also predicts a few rare variants as P/LP in these genes which have not been reported previously in ClinVar. Investigators who intend to use the ExAC noTCGA dataset as controls in cancer sequencing studies can use PathoMAN to get a rapid count of variants across genes above and beyond those reported in ClinVar.

**Table 4:**
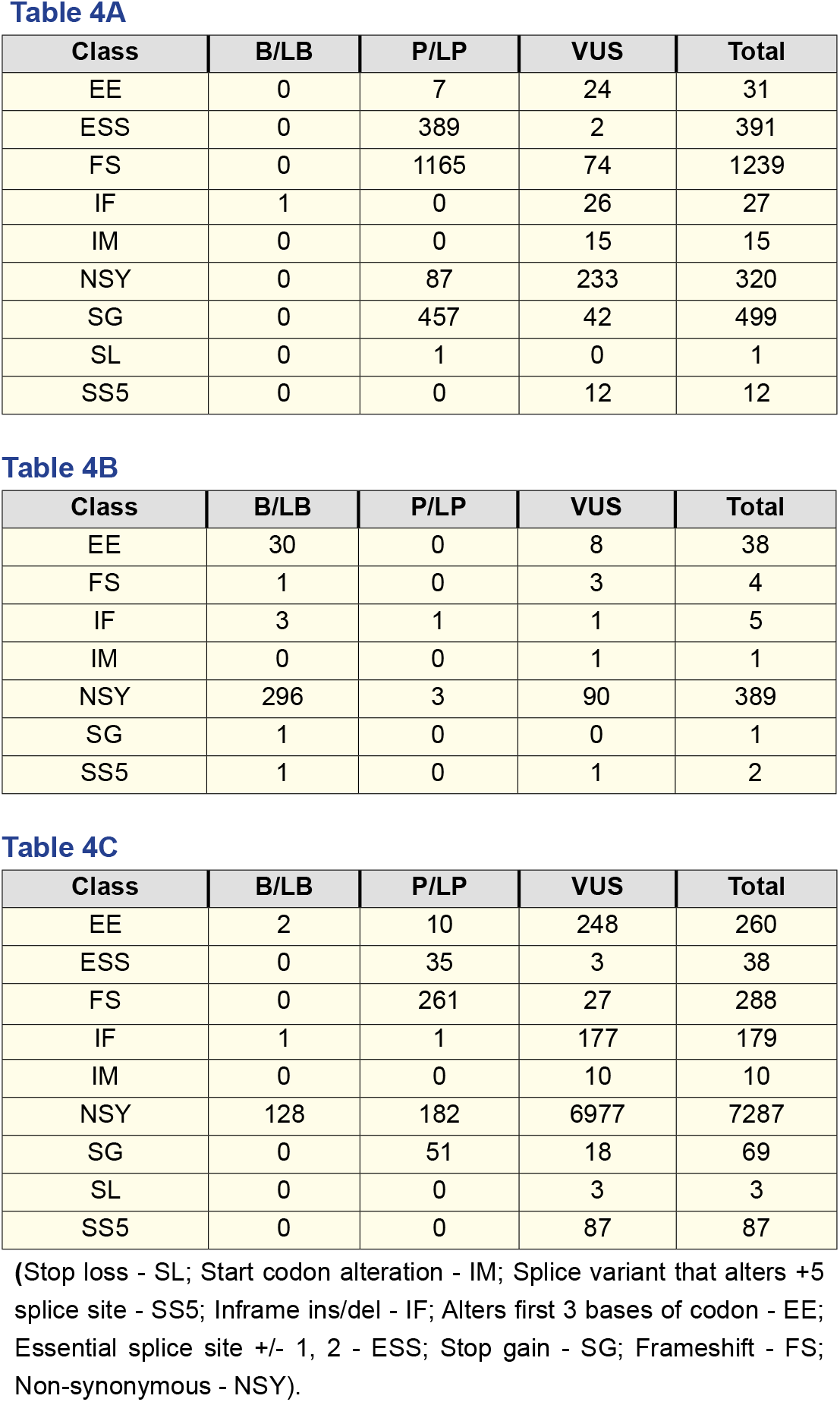
PathoMAN re-classification of expertly curated germline cancer variant reported in the test datasets. All tables show distribution by variant classes. **Table A** shows only reported P/LP variants; **Table B** shows only reported B/LB; and **Table C** shows only Reported VUS.

| Class | B/LB | P/LP | VUS | Total |
| --- | --- | --- | --- | --- |
| EE | 0 | 7 | 24 | 31 |
| ESS | 0 | 389 | 2 | 391 |
| FS | 0 | 1165 | 74 | 1239 |
| IF | 1 | 0 | 26 | 27 |
| IM | 0 | 0 | 15 | 15 |
| NSY | 0 | 87 | 233 | 320 |
| SG | 0 | 457 | 42 | 499 |
| SL | 0 | 1 | 0 | 1 |
| SS5 | 0 | 0 | 12 | 12 |

| Class | B/LB | P/LP | VUS | Total |
| --- | --- | --- | --- | --- |
| EE | 30 | 0 | 8 | 38 |
| FS | 1 | 0 | 3 | 4 |
| IF | 3 | 1 | 1 | 5 |
| IM | 0 | 0 | 1 | 1 |
| NSY | 296 | 3 | 90 | 389 |
| SG | 1 | 0 | 0 | 1 |
| SS5 | 1 | 0 | 1 | 2 |

| Class | B/LB | P/LP | VUS | Total |
| --- | --- | --- | --- | --- |
| EE | 2 | 10 | 248 | 260 |
| ESS | 0 | 35 | 3 | 38 |
| FS | 0 | 261 | 27 | 288 |
| IF | 1 | 1 | 177 | 179 |
| IM | 0 | 0 | 10 | 10 |
| NSY | 128 | 182 | 6977 | 7287 |
| SG | 0 | 51 | 18 | 69 |
| SL | 0 | 0 | 3 | 3 |
| SS5 | 0 | 0 | 87 | 87 |

(Stop loss - SL; Start codon alteration - IM; Splice variant that alters +5 splice site - SS5; Inframe ins/del - IF; Alters first 3 bases of codon - EE; Essential splice site +/- 1, 2 - ESS; Stop gain - SG; Frameshift - FS; Non-synonymous - NSY).

### Usage of ACMG/AMP categories in PathoMAN

We analyzed the real-world usage of the eight categories of evidence (population frequency, genomic annotation and computational prediction, functional evidence, co-segregation, *de novo* status, allelic/genotypic data, public databases, scientific literature and other data) used in the ACMG/AMP guidelines. Interestingly, we find that the categories: population frequency data, genomic annotation and computational predictions, databases and scientific literature (**Figure 2**) are the most used. These are available due to generous data and tool-kit sharing policies in the genomics field. The categories that are rarely if ever used are familial cosegregation data or *de novo* status, allelic data, and functional data. The co-segregation data and *de novo* status data are limited to familial studies, and are mostly unavailable in sporadic case-control settings since these are collected by investigators, doctors, genetic counsellors and commercial labs based on patient input. For a variant to be classified as pathogenic or likely pathogenic by ACMG criteria, one needs a maximum of 1 PVS1 or 2 PSs or 3 PMs or 4 PPs for which, the knowledge-base and resources used by PathoMAN were demonstrably sufficient. We describe below the bottlenecks in sharing this information and propose a novel framework to circumvent and ameliorate these issues.

**Figure 2:**
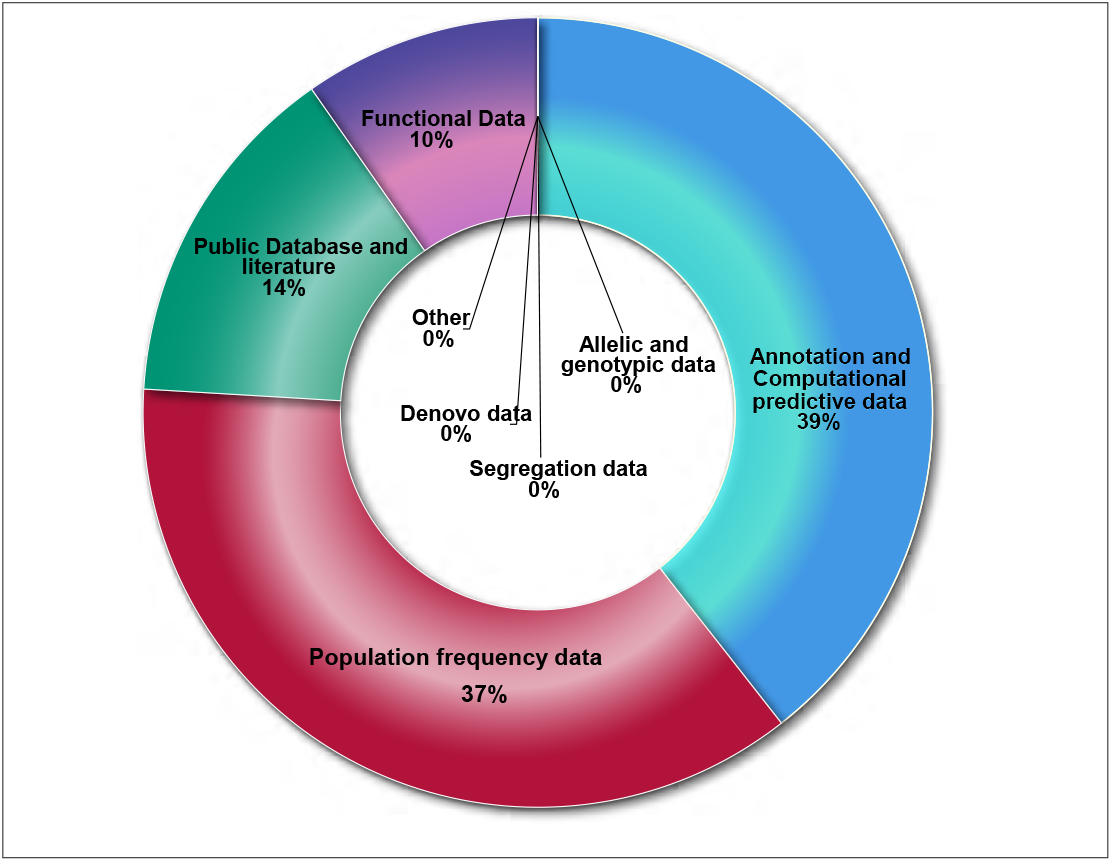
Utilization of knowledgebase components by PathoMAN during variant curation of the test datasets.

**Table 5:** Comparison of pathogenic gene burden in ExACnoTC-GA between ClinVar and PathoMAN. Columns contain variant counts.

| GENE | ClinVar P/LP | PathoMAN P/LP | Difference |
| --- | --- | --- | --- |
| *BLM* | 1 | 9 | 8 |
| *TP53* | 15 | 7 | 8 |
| *ATM* | 80 | 74 | 6 |
| *BRCA2* | 101 | 96 | 5 |
| *BARD1* | 10 | 15 | 5 |
| *SDHA* | 5 | 0 | 5 |
| *MLH1* | 6 | 11 | 5 |
| *EGFR* | 0 | 5 | 5 |
| *RAD51B* | 0 | 5 | 5 |
| *PALB2* | 21 | 26 | 5 |
| *RAD50* | 21 | 17 | 4 |
| *PMS2* | 15 | 11 | 4 |
| *CDH1* | 3 | 7 | 4 |
| *MRE11A* | 11 | 7 | 4 |
| *EPCAM* | 0 | 4 | 4 |
| *BRCA1* | 68 | 65 | 3 |
| *NF1* | 3 | 6 | 3 |
| *PTEN* | 2 | 5 | 3 |
| *STK11* | 0 | 3 | 3 |
| *KRAS* | 2 | 0 | 2 |
| *MUTYH* | 26 | 24 | 2 |
| *BRIP1* | 21 | 23 | 2 |
| *RAD51C* | 17 | 15 | 2 |
| *FH* | 6 | 4 | 2 |
| *RET* | 2 | 4 | 2 |
| *BMPR1A* | 1 | 3 | 2 |
| *FAM175A* | 1 | 0 | 1 |
| *RAD51* | 1 | 0 | 1 |
| *MSH6* | 13 | 12 | 1 |
| *NBN* | 10 | 9 | 1 |
| *RAD51D* | 6 | 7 | 1 |
| *APC* | 4 | 5 | 1 |
| *CDKN2A* | 5 | 4 | 1 |
| *DICER1* | 3 | 2 | 1 |
| *BAP1* | 0 | 1 | 1 |

## Discussion

### PathoMAN as a tool to aid variant curation

Traditionally, genetic variant curation has been performed manually by expert groups of individuals. However this is a time intensive task that requires aggregation and interpretation of information from multiple sources. In the cancer realm, this was relatively easy at times when only a single gene such as *BRCA1/2* was under investigation. In contemporary testing scenarios which routinely rely on multiplex gene-panels, this task is onerous. Large gene discovery efforts, as well as clinical reporting, could use a simplified, automated, method for prioritizing variants for a closer look or in the best case, be useful as the classification tool of choice. PathoMAN addresses this critical unmet need for an unbiased algorithmic approach towards classifying genetic variants of clinical interest in cancer predisposition. PathoMAN can be easily accessed through a web browser and results for individual variants are almost immediately available, while batch uploads of variant VCFs may take a few hours.

Genetic testing labs have started utilizing the ACMG/AMP classification rules to classify variants for pathogenicity within cancer predisposition genes. However, results vary depending on availability of accessible data and interpretational differences^55^. Efforts are being made to resolve interpretational differences through initiatives underway such as ClinGen. In a recent report^56^, 13% of variants in ClinVar were re-analyzed, and were found to be unresolved, underscoring the difficulties even for expert curator groups. In that study^56^, clinical intepretations from four clinical laboratories were concordant for 91.5% of shared variants in ClinVar after consulations. For manual or automated curation the minimal set of information required to classify a variant as likely pathogenic or likely benign are: population frequency, computational predictors and evidences from public databases^56^. PathoMAN compiles this information uniformly in a machine accessible format which is used as a knowledge-base for variant classification. An advantage of using PathoMAN is that it can easily tag benign variants based on public allele frequency and the genomic context information from annotators. This reduces the variant pool of interest to a manageable subset. In a typical multiplexed gene-panel variant list, after filtering for only rare high or moderate impact variants, PathoMAN will classify about one third of the variants as B/LB with high precision. This saves time and effort for the variant curators and helps them to focus on curating the remaining potentially actionable variants. PathoMAN can also tag founder mutations.

Cancer is a complex disease with multi-gene aetiology. Some cancer genes confer high risk whereas some only moderately affect the carrier’s risk. Panel testing is currently used for active surveillance and intervention to lower disease risk. Large sequencing and genotyping efforts to discover new cancer predisposition genes are being carried out by several consortia like BCAC^57^, SIMPLEXO^58;^ COMPLEXO^59^, CIMBA^60^, etc. Several commercial and academic labs also now offer multiplexed panels. As the cost for sequencing these panels decreases, the number of genes tested in panels is increasing. Automation allows for rapid processing, service assurance and reproducibility of results for these large panels. Gold standard sets of curation pioneered by ClinGen^51; 52^ would aid in refining these pathogenicity classifications further, while efforts such as the PROMPT^29^ registry enable accurate penetrance estimates of mutations in susceptibility genes. The PROMPT registry has identified a 26% discordance rate among laboratories and an 11% rate with conflicting interpretations, a discrepancy that has implications for altering medical management.

In the three datasets we used to test PathoMAN, we demonstrate that, when contrasted against an expert curated set of variants in the IMPACT-76 genes, there is a high concordance rate for both pathogenic and benign variants. The concordance for P/LP variants is excellent when limited to truncating variants; frameshift variants and essential splice variants such as those reported in the prostate cancer study^8^. When missense variants are also considered such as in the breast cancer study^9^, we see more VUS, but the discordance is still minimal.

Much of the discordance can be traced back to the ClinVar submissions with conflicting interpretations; e.g. *BRCA1*:c.5348T>C (p.Met1783Thr). Currently, PathoMAN doesn’t have access to proprietary databases (such as HGMD)^62,63^, which may have additional evidence for pathogenicity or benignity. For splice variants, our source is limited to −3-to+8 at the 5’ splice site and −12-to+2 at the 3’ splice site. Hence, we may be missing out on certain extended regions.

Many labs and certain programs such as cardio classifier^64^ and InterVar^65^ use prior knowledge of disease-gene pair association. This is advantageous to reduce classifications leading to P/LP for those genes that are not in a disease-gene pair. However, it also suffers from the disadvantage that it cannot be used for lesser known genes-disease pairs or for novel gene hunting. In a recent report, we showed that, half of the cases, in a series consisting of selected advanced cancers at a single institution, were non-syndromic associations^5^. Probands or their close relatives had clinically actionable variants in cancer genes not directly associated with the specific cancers for which there were known syndromic associations. PathoMAN does not use the contextual syndromic association in deciphering pathogenicity of variants. However, with the applications envisaged for novel gene discovery, this is a distinct advantage. For clinical sequencing which is more focused on specific sets of genes, is limiting to disease-gene pairs to identify pathogenic variants.

The variants that could not be classified by PathoMAN and are called VUS are due to lack of accessible, supporting evidence for the clinical assertion by using ACMG guidelines. These are classic examples of rare variants absent in ClinVar and ExAC datasets. For these LOR variants, we believe that the expert curators may have had additional evidence form literature, inhouse functional evidence^66^ or familial co-segregation information^67^ that helped classify these variants as P/LP or B/LB. The upgrade for VUS to either LP or LB by PathoMAN is based on the three categories - lines of available evidence in public databases, population frequency and computational and in silico prediction on deleteriousness. These variants can be re-classified as either pathogenic or benign if additional functional or co-segregation data become available through literature or initiatives such as ClinGen^56^.

Commercial testing laboratories have proprietary versions of interpretation pipelines such as Sherloc68(Invitae Corporation) and MyVISION (Myriad Genetics). However, these are unavailable to the community at large. PathoMAN is designed to provide an optimized platform for clinical variant calling utilizing publically available data resources.

### Using ACMG for variant classification in Cancer

Mutations in tumor suppressors and oncogenes lead to tumorigenesis, and the Knudson two-hit hypothesis69is seen to operate in many common cancers. Common examples include *APC, TP53, BRCA1/2* genes etc. However, several of these genes, especially those that are part of the Fanconi complex (*FANCS-BRCA1*, *FANCD1-BRCA2*, *FANCJ-BRIP1*, *FANCN-PALB2*, *FANCP-SLX4*, *RAD51C*), neurofibromatosis (NF1), Ataxia-telangiectasia (*ATM*), Bloom syndrome (BLM), Niemegen breakage syndrome (NBN), dyskeratosis congenita (*TERT*) that lead to autosomal recessive rare Mendelian disorders, are also found to be risk genes for autosomal dominant cancer predisposition. Heterozygous carriers of these gene mutations are reported to have increased risks for syndromic cancers^70^. Occasionally, gene disrupting heterozygous mutations in these genes that are rare, absent in public controls such as ExAC and gNOMAD may be observed in sequenced cancer cohorts. Their ClinVar record for pathogenicity is usually based on their Mendelian recessive syndrome and not to the cancer phenotypes. Hence, applying the ACMG rules to genes without membership in the ACMG list may be fraught with misclassification. However, we believe that continuing data streams for variants in these genes will lead to better classifications, especially when coupled with familial co-segregation and functional validations. While PathoMAN classifications for such genes are a useful starting point for identifying variants that may be pathogenic, and discarding benign; we emphasise on expert manual curation to disentangle these issue.

### Limitations of automation

Automating variant classification based on publically available information has some pitfalls. Supporting evidences provided in ClinVar for variants are not computation friendly and requires manual curation to interpret free text. In several instances, the citations are not relevant to the specific records. Technologies such as natural language processing and tagging will eventually help to build a knowledge-base that can further be used for deep learning.

Current ACMG guidelines do not directly link ClinVar functional evidence provided as supporting observations, which leads to loss of information that could be used in variant classification. Due to this lack of data structure, the variants in ClinVar are scored only PP5 or BP6 and not PS3 or BS3. We employed the gold star 2 or more status as a proxy for functional evidence. Not all clinical sequencing projects are equipped or do independent analyses to assess functional evidences for their clinical assertion. If the ClinVar evidence is coded with proper tags, it would be helpful for molecular geneticists and clinical curators to use this information for their pathogenicity estimation. For example *TP53* (R273H), *BRCA1* (Y105C) and *BRCA1* (V1688del) variants have overwhelming literature evidences (**Supp Figure 2**); however the evidence present in the description of the submissions within ClinVar, are computationally un-derivable. Similarly there are many variants reported in the literature which may have some level of supporting evidence for pathogenicity or benignity in ClinVar. Currently all of these data integration is done by manual curators on a case-by-case basis.

We propose a framework to report ClinVar data that can be structured and parsable for an automated algorithm in the context of cancer. This format consists of 6 important fields that compress the vast information that is present in literature or clinical reports.

1. Population/Ethnicity (NFE, AFR, SAS, AMR, ASJ, FIN, OTH, EAS, others)
2. Inheritance model (AD,AR, *de novo*, X-linked)
3. Allelic status (Hom, Het)
4. Family history/Co-segregation information (Yes-1; No-0)
5. Disease association (TCGA code/ Oncotree code^71^)
6. Functional Evidence (Experiment type: NMC, LOH, etc.)

For example, *ERCC3* (R109X) variant^72^ can be depicted as ASJ-AD-Het-1:1-BRCA,BLCA-NMD. This variant was seen in Ashkenazi Jewish individuals with an autosomal dominant inheritance for the heterozygous allele. This variant cosegregated in one family with cancer history. The variant was found in Breast cancer and Bladder cancer individuals and the functional evidence for pathogenicity was carried out by testing for non-sense mediated decay and other experiments.

Large sequencing studies and gene specific functional studies give curated list of variants with their pathogenic impacts like TP53 database^28^ and a functional study on *PALB2* variants^26; 27^. As a primer, we have collated a list of *PALB2, TP53* variants from the literature as supporting the knowledge-base for PathoMAN but there is a real need to create a publically available well curated list of variants from the literature that is amenable to programmatic interpretation. Similarly, as standards evolve for the incorporation of somatic mutations into germline interpretation, we expect an integration of such events for atleast some tumor suppressor and oncogenes. The roles played by the ENIGMA Consortium^73; 74^, G4GH^75^, BRCA-Share^76^ in this regard are meritorious. Though Clinical laboratories collaborate to resolve the differences in variant interpretations submitted to ClinVar^56^, the fact remains however, that a unified framework for incorporation of supporting machine readable evidences in any variant database including ClinVar remains a critical bottleneck.

Functional data is rarely available for most genes. Exceptions are *BRCA1/2* due to the concerted efforts of the ENIGMA consortium^73; 74^. In single variant reports, data is usually buried within scientific jargon that is not compatible with genomic variant information. In many instances, functional data is dependent on the models used, e.g. overexpression of a mutant construct, deletion of a region using a CRISPR endonuclease and sometimes, introduction of the specific nucleotide through homology directed DNA repair. It is also likely, that the results from these three methods do not agree. Novel methods to understand deleteriousness using saturation mutagenesis are also starting to emerge^77–79^ for e.g., BRCA1^80^ that we have incorporated into PathoMAN. We hope these will add a uniform layer of functional data that can be used in determining pathogenicity in the coming years.

In conclusion, we performed pathogenicity assessment of 66,762 variants in germline cancer genes (IMPACT-76), the first and largest uniform classification using an unbiased computational tool. We demonstrate the high concordance and low discordance when compared with manual curation as a harbinger of how such programs will in the near future, be able to work as well as domain experts and manual curators. PathoMAN is a first step towards our goal of automating the complex process of variant classification and interpretation. A beta version of the web app is available at https://pathoman.mskcc.org/

## Web -resources

1. CAVA - https://github.com/RahmanTeam/CAVA
2. SNPEff - http://snpeff.sourceforge.net/SnpEff_manual.html
3. Annovar - https://annovar.openbioinformatics.org
4. ExACnoTCGA - http://exac.broadinstitute.org
5. gnomAD - http://gnomad.broadinstitute.org/
6. ClinVar - https://www.ncbi.nlm.nih.gov/clinvar/
7. IARC database - http://p53.iarc.fr/
8. ClinVar parser tool - https://github.com/macarthur-lab/clinvar
9. dbNSfP and dbscSNV - https://sites.google.com/site/jpopgen/dbNSFP
10. Gene List - https://github.com/macarthur-lab/gene_lists
11. Repeat masker - http://www.repeatmasker.org/
12. UCSC Genome Browser - https://genome.ucsc.edu
13. Cardio classifier - https://www.cardioclassifier.org/
14. InterVar - https://github.com/WGLab/InterVar
15. ACMG - https://www.acmg.net/
16. PathoMAN- http://pathoman.mskcc.org/

## Acknowledgements

We thank Sabine Topka PhD, Semanti Mukherjee PhD, Maria Carlo MD and Zoe Steinsynder BS for helpful suggestions to improve the manuscript.

## Funding/Support

Research reported in this pre-print was supported by National Cancer Insittute of the National Institutes of Health under award number R21CA029533 and R21CA178800 as well as Cycle for Survival, the Breast Cancer Research Foundation and The V Foundation for Cancer Research. It is also supported by the Cancer Center core grant P30CA008748 and The Robert and Kate Niehaus Center for Inherited Cancer Genomics. The content is solely the responsibility of the authors and does not necessarily represent the official views of the National Institutes of Health or other funding agencies.

## Supplementary Information

**Supplementary Table 1:**
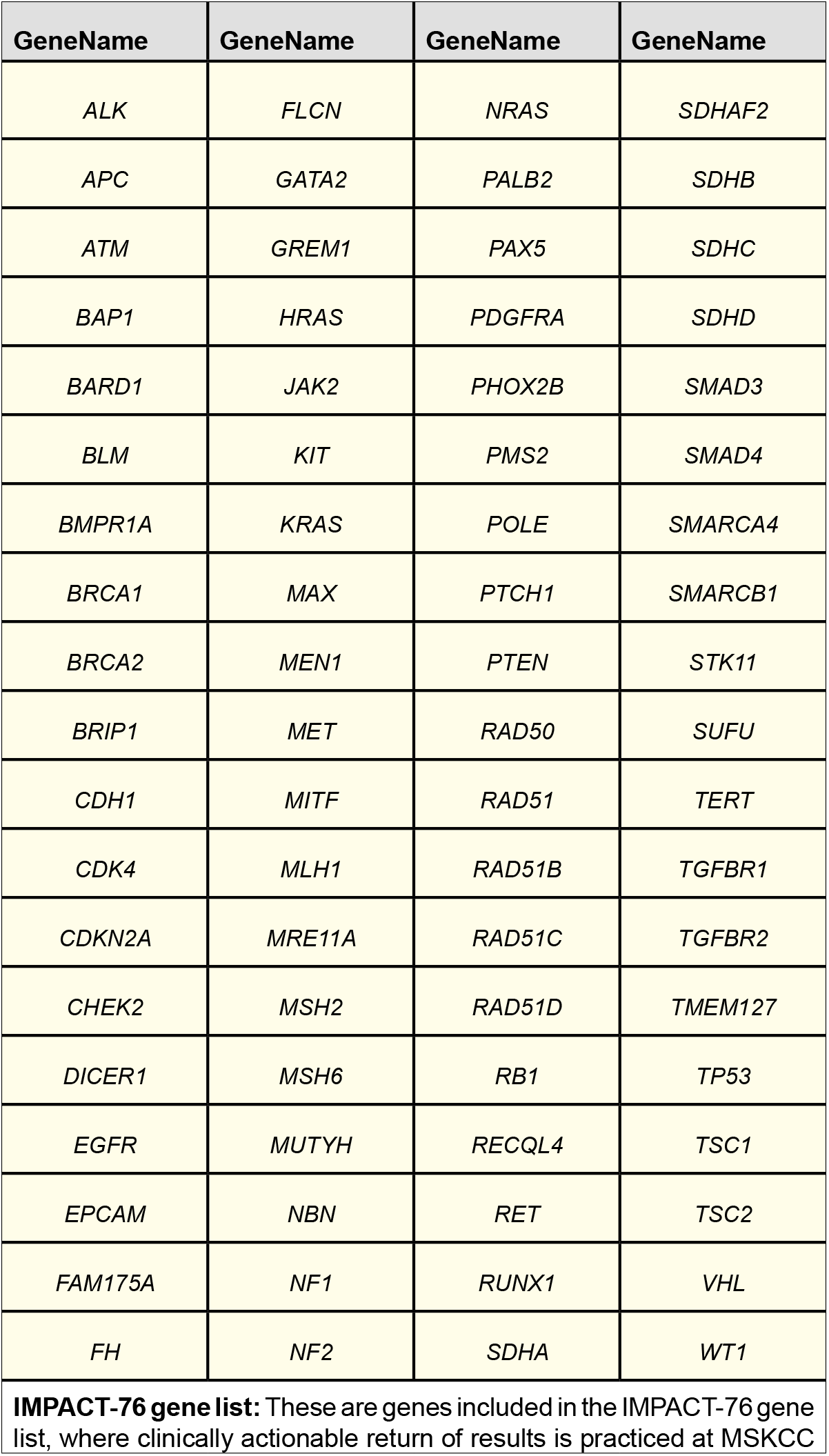

| GeneName | GeneName | GeneName | GeneName |
| --- | --- | --- | --- |
| *ALK* | *FLCN* | *NRAS* | *SDHAF2* |
| *APC* | *GATA2* | *PALB2* | *SDHB* |
| *ATM* | *GREM1* | *PAX5* | *SDHC* |
| *BAP1* | *HRAS* | *PDGFRA* | *SDHD* |
| *BARD1* | *JAK2* | *PHOX2B* | *SMAD3* |
| *BLM* | *KIT* | *PMS2* | *SMAD4* |
| *BMPR1A* | *KRAS* | *POLE* | *SMARCA4* |
| *BRCA1* | *MAX* | *PTCH1* | *SMARCB1* |
| *BRCA2* | *MEN1* | *PTEN* | *STK11* |
| *BRIP1* | *MET* | *RAD50* | *SUFU* |
| *CDH1* | *MITF* | *RAD51* | *TERT* |
| *CDK4* | *MLH1* | *RAD51B* | *TGFBR1* |
| *CDKN2A* | *MRE11A* | *RAD51C* | *TGFBR2* |
| *CHEK2* | *MSH2* | *RAD51D* | *TMEM127* |
| *DICER1* | *MSH6* | *RB1* | *TP53* |
| *EGFR* | *MUTYH* | *RECQL4* | *TSC1* |
| *EPCAM* | *NBN* | *RET* | *TSC2* |
| *FAM175A* | *NF1* | *RUNX1* | *VHL* |
| *FH* | *NF2* | *SDHA* | *WT1* |

**IMPACT-76 gene list:** These are genes included in the IMPACT-76 gene list, where clinically actionable return of results is practiced at MSKCC

**Supplementary Figure 1:**
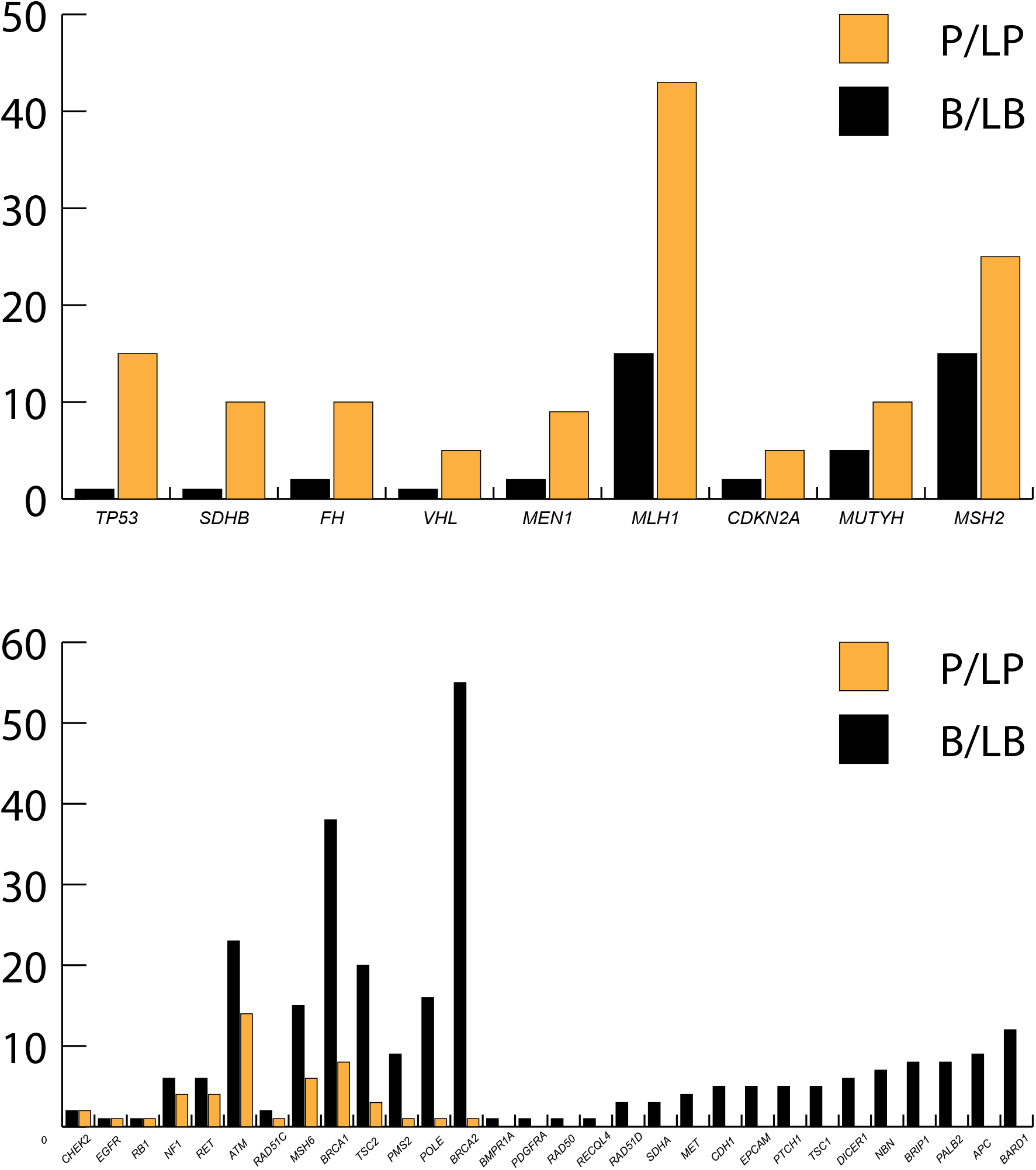
(Top): Histogram of cancer related genes with high ratio of pathogenic missense variants from ClinVar; (Bottom): Histogram of cancer related genes with high ratio of benign missense variants from ClinVar.

**Supplementary Figure 2.**
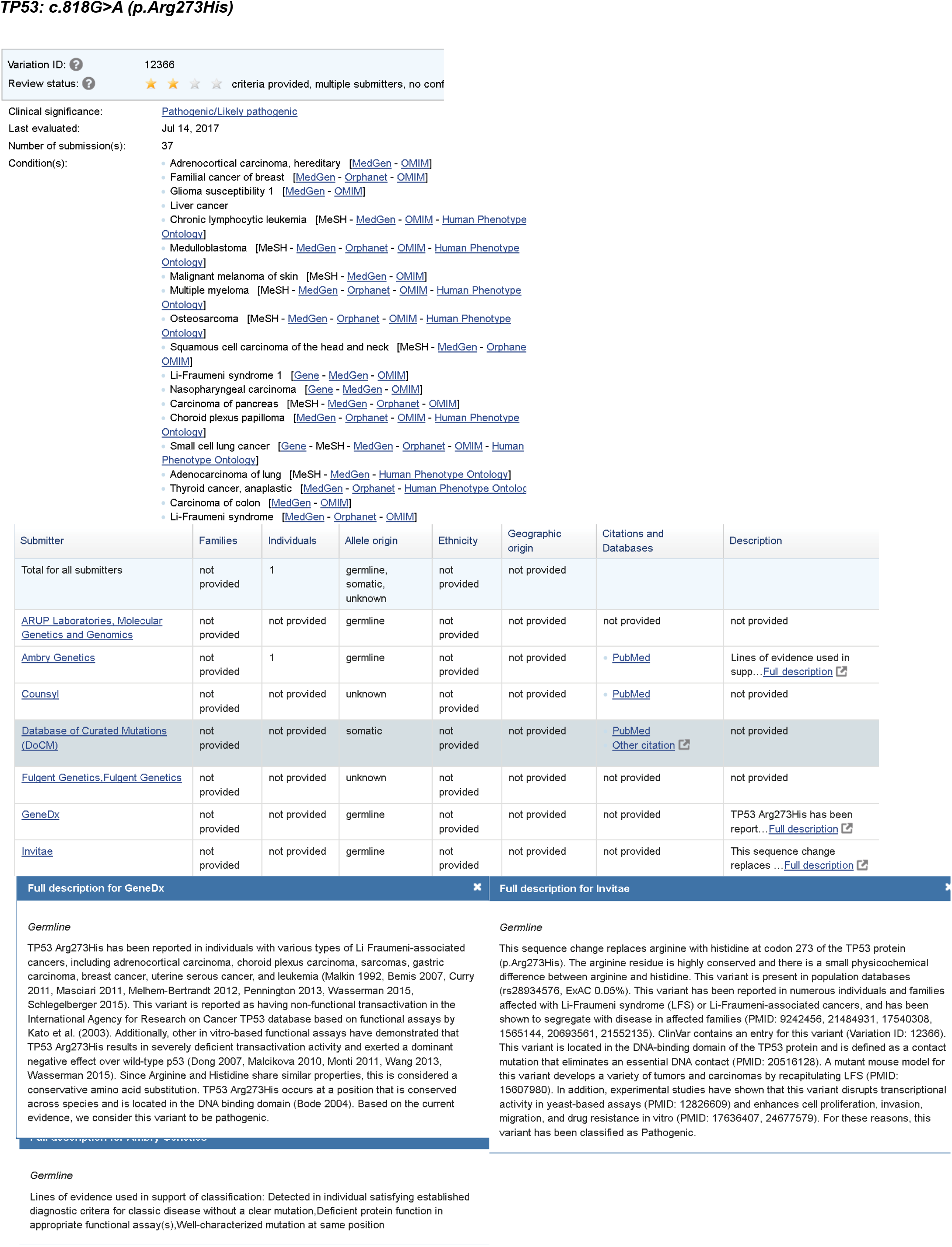
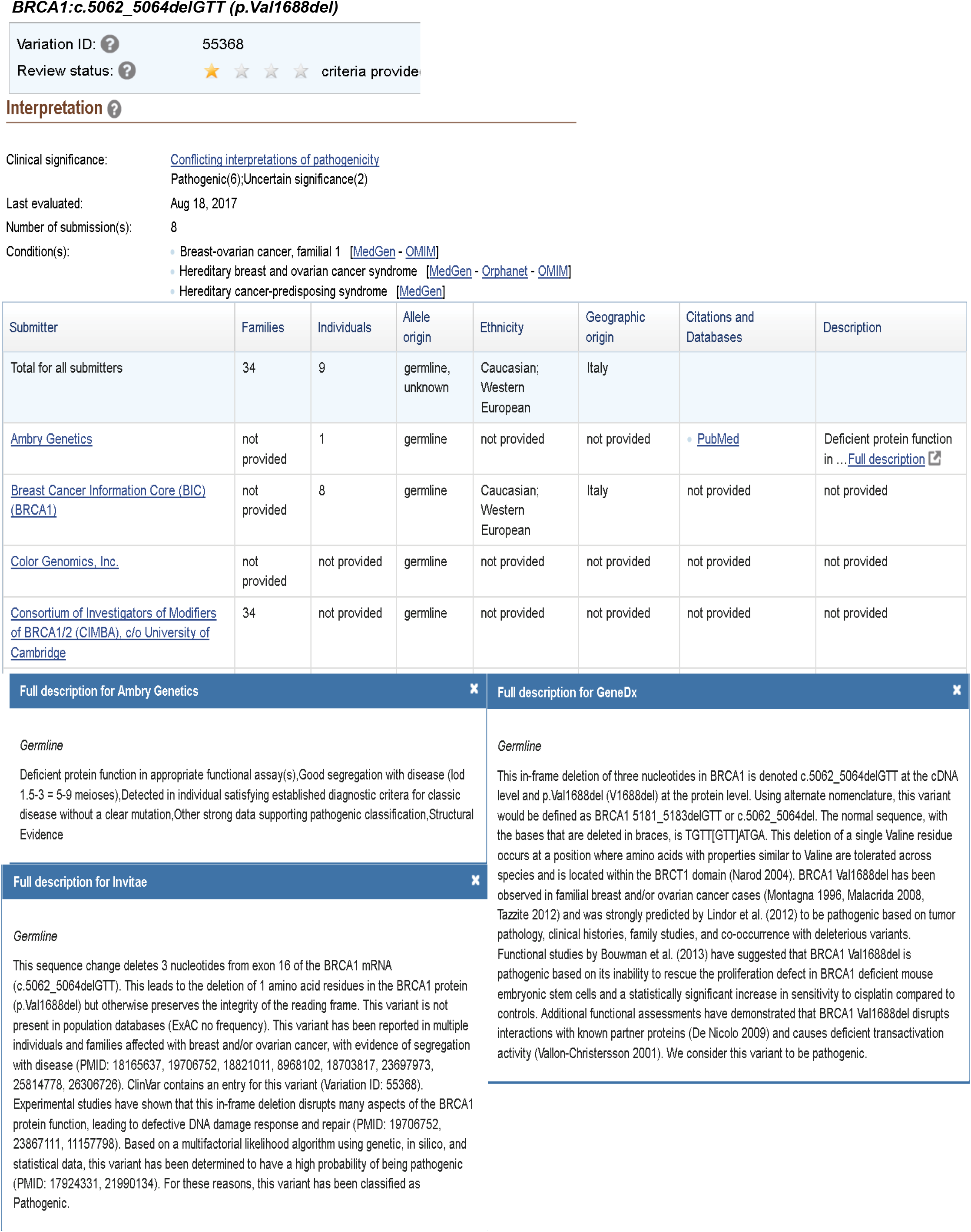
TP53: c.818G>A (p.Arg273His)

